# Vision guides directed orienting movements during obstacle avoidance in mice

**DOI:** 10.64898/2026.03.12.711382

**Authors:** Michael Sidikpramana, Keaton Jones, Cristopher M. Niell

## Abstract

In natural environments, animals must effectively maneuver around obstacles to reach goals such as food or shelter. Recent work has demonstrated that laboratory mice use vision for naturalistic behavior such as prey capture, escape, and distance estimation. However, it is unknown to what extent mice use vision relative to other senses such as touch for obstacle avoidance, a critical natural behavior. In this study we developed an obstacle avoidance task in freely moving mice to investigate how vision is used to guide paths around an obstacle obstructing a goal. We found that mice clearly use vision to avoid an obstacle, steering around the obstacle at distances where tactile information isn’t available. By comparing trajectories for mice performing obstacle avoidance in the light versus the dark, we found that vision contributes to more spatially efficient trajectories and paths directed to the open edge of the obstacle. When vision is available, mice make large orienting movements towards the opening of the obstacle at about 10 cm from its edge, suggesting that mice are actively using visual information to direct these movements. Finally, by occluding one eye, we found that mice were still able to avoid obstacles with primarily monocular information. Taken together, these results demonstrate that laboratory mice use vision to avoid an obstacle, taking directed paths that are initiated by large orienting movements. In addition to demonstrating the visual behavioral capabilities of the mouse, this paradigm can serve as a foundation to study the neural circuits that mediate visually guided orienting and locomotion.

**Highlights:**

- We developed a simple obstacle avoidance task for freely moving mice that requires minimal training
- Vision is necessary for efficient and directed paths around an obstacle
- Mice steer around obstacles by performing directed head movements towards clear paths
- Mice do not require binocular vision for obstacle avoidance

## Introduction

In nature, vision is an incredibly active process - animals shift their gaze across the visual scene, identifying relevant objects in the environment with different behavioral contingencies, such as food, predators, and obstacles. Researchers have used ethologically relevant visual behavior in rodents to study the active interplay between visual information and motor action. For example laboratory mice use vision to track and capture insects, identify and escape from threatening stimuli, and judge gap distances for jumping^1,2,3,4,5^. Despite having low acuity vision, mice are a powerful system to study natural vision, due to their rich repertoire of visual behaviors, similarities of the visual system to other mammals, and genetic access ^6,7^.

One of the primary uses of vision is to navigate around obstacles in complex environments, a prevalent behavior across the animal kingdom^8,9,10^. In predictable environments such as our own homes, we often rely on spatial representations that are learned with sensory information such as vision, but rely less on sensory input once committed to memory. For example, a recent study showed that when learning to avoid a stationary/predictable obstacle during an escape task, mice primarily used spatial memory rather than vision, memorizing the location of the obstacle over the course of an experiment^11^. When the obstacle appeared at stimulus onset or unexpectedly, mice could use vision but relied on primarily tactile cues to direct their paths around the obstacle^11^. This aligns with other escape literature which suggest that spatial representations present in the retrospinal cortex or hippocampus are used for path planning during escape/instinctive behaviors^12^. In nature, environments are dynamic and unexpected obstacles may be encountered that are not committed to memory yet. It is still unknown if mice use vision to avoid obstacles at a distance, similar to orienting during approach/prey capture behavior, or if mice rely on other senses like touch, known to drive other directed locomotion^13,14^. In human studies of obstacle avoidance and navigation, studies have shown that the behavioral dynamics of steering were reflective of active visual control^15,16^. In particular gaze duration and sequence across the obstacle, goals, and current path reflect the need of online visual information during steering. It is not clear to what extent these dynamics are reflected in mice, especially since mice are non-foveated, have near panoramic visual fields, and change their gaze primarily by leading with their heads^17,18^.

To study whether mice can use active vision to guide steering in an obstructed environment, we developed a naturalistic task that taps into the innate behavior of obstacle avoidance. Our experiments clearly showed the use of vision to avoid an unexpected obstacle. By comparing paths and steering in light and dark conditions, we determined the behavioral dynamics of visually guided obstacle avoidance. Visually driven paths were more directed at the edge of the obstacle and were initiated by large and accurate orienting movements towards the opening next to the obstacle. In contrast, in the dark, mice continued on their initial trajectory and would often hit the obstacle. By occluding a single eye, we found that mice had no deficits when the obstacle was on the non-occluded side and could still accurately target the obstacle opening, whereas paths were slightly less directed when the obstacle opening was on the side of the occluded eye. This paradigm gives insights into how mice use vision to guide locomotion around an obstacle and provides a foundation to study the corresponding neural computations.

## Results

### Freely moving obstacle avoidance task

In order to study how mice steer around an obstacle, we first trained mice to collect water at ports at each end of an arena (Fig 1A,C). After a water reward delivery at one port was triggered by nose poke, the port at the other end of the arena became active, thus requiring mice to traverse back and forth across the arena. We define a single trial as the time to traverse the arena from one water port to another. Within 3-5 days mice reached an average of 111 ± 10 trials per session and 4 ± 0.30 seconds per trial (Fig 1E, S1A). This benchmark was sufficient to receive adequate water (∼2-3ml daily) within sessions of approximately 15 mins. We then capped the session duration of obstacle experiments to a maximum of 15 mins. Sessions ended early if mice were not collecting water to encourage continuous water collection throughout a session. After reaching these benchmarks, an opaque wall-like obstacle was introduced into the arena, which mice had to locomote around while collecting water (Fig 1C). During the obstacle avoidance experiments, the obstacle was moved randomly by hand to one of six positions every three trials, therefore mice would see the obstacle at the same position at the same distance twice before switching locations (Fig 1D). We found that there was relatively little difference in behavior depending on repeated presentations of the obstacle, suggesting that mice may not use memory to perform this task (S5 A-E). After 7 days of performing the basic obstacle avoidance task, we tested different manipulations for 3-5 days, which included darkness or monocular occlusions (Fig 1B). During all sessions, overhead video was acquired using a 60 fps camera (Flir Blackfly usb3) and markerless pose estimation was performed post hoc using DeepLabCut (Fig 1A)^19^.

**Figure 1.**
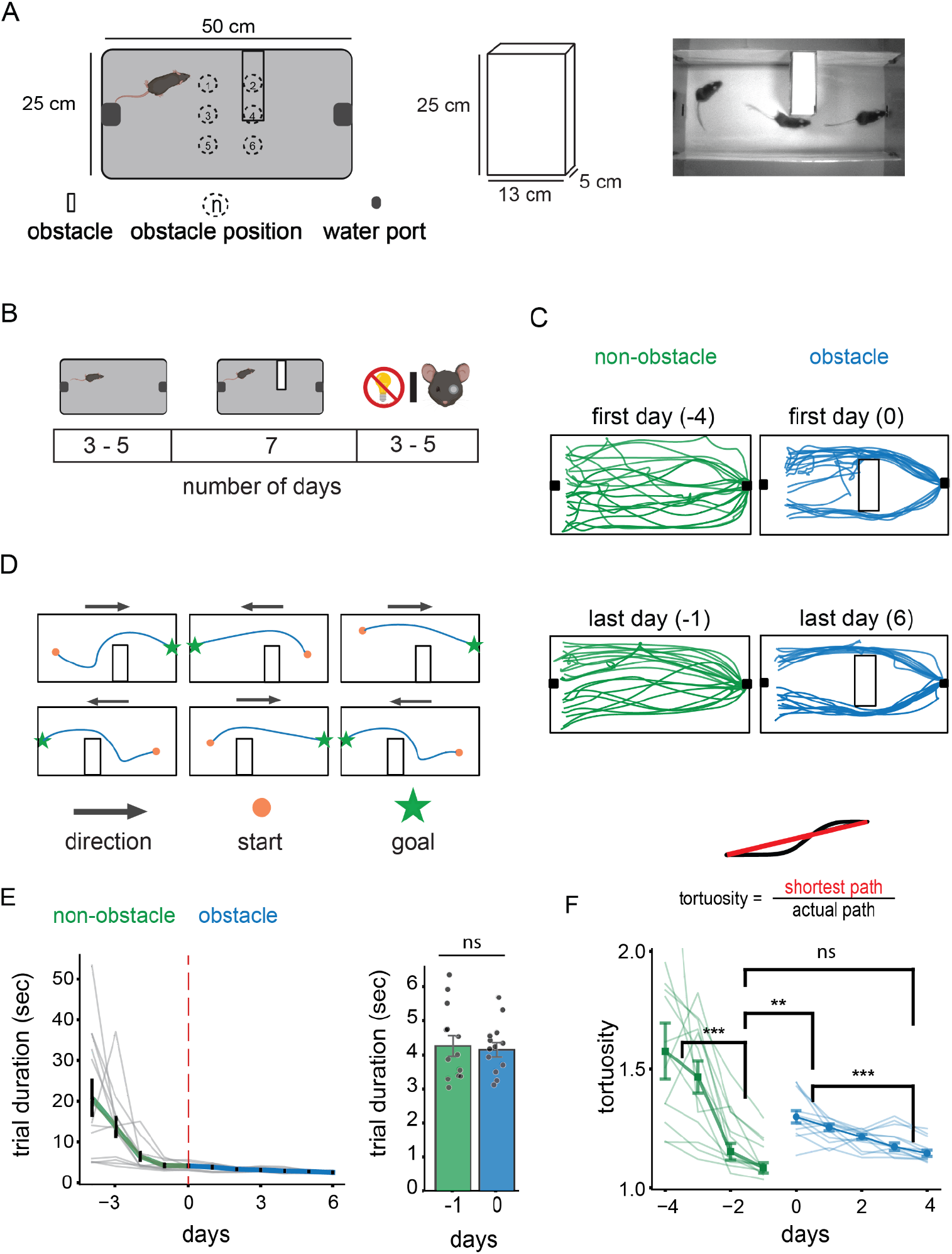
Obstacle avoidance task design and training. (A) Left, schematic of obstacle avoidance arena. Middle, schematic of wall-like obstacle. Right, overlaid example frames from a single trial. (B) Experimental timeline: Left, non-obstacle 3-5 days. Middle, obstacle task 7 days. Right, task manipulation 3-5 days, darkness or monocular occlusion. (C) 20 representative trials from the first and last day without (left) and with (right) the obstacle. (D) Schematic of obstacle avoidance task structure and example trajectories. (E) Left, between animal and single animal mean trial duration across days during non-obstacle and obstacle phase. Right, mean trial duration of the last day of non-obstacle and first day of obstacle task. (F) Between animal and single animal median tortuosity across days during non-obstacle and obstacle phase. Above: Tortuosity schematic. One outlier point not shown on day -4 omitted for visualization purposes because it exceeds the y-axis scale. N = 13 mice.

To compare how spatially efficient paths were in each condition, we calculated the tortuosity, which is the ratio between the shortest possible path and actual path (Fig 1F). A two-way ANOVA showed an effect of condition (non-obstacle vs obstacle) and day but not an interaction between the two factors (non-obstacle vs obstacle: p = 0.0001, days: p = 0.004, interaction: p = 0.601). We found that tortuosity was significantly less on late non-obstacle sessions (days -2,-1) relative to early obstacle sessions (days 0,1) (Wilcoxon signed rank; p=0.0061, non-obstacle late: 1.11 ± 0.04, obstacle early: 1.27 ± 0.02), but no significant difference when comparing late non-obstacle with late obstacle sessions (Wilcoxon signed rank; p= 0.4, obstacle late: 1.16 ± 0.02) (Fig 1F). We also found a significant decrease in tortuosity within conditions from early to late sessions, demonstrating that mice continually improved their spatial efficiency across days (non-obstacle; early: 1.54 ± 0.12, late: 1.13 ± 0.04, Wilcoxon signed rank; p< 0.001, obstacle; early: 1.27 ± 0.02, late: 1.16 ± 0.02, Wilcoxon signed rank; p< 0.001). These data show that mice require little training for our obstacle avoidance task and demonstrate that they effectively traverse the arena to collect water with an obstacle present. This is consistent with obstacle avoidance as a natural behavior that would be performed constantly during movement through the environment in the wild.

Depending on the mouse’s initial position at the start of the trial, the obstacle may not obstruct the path to the reward port, thus not requiring active avoidance. Therefore we focused our analysis on trials in which the obstacle was in a lateral position in the way of the mouse’s path to the water port. Additionally, when the obstacle was at the closer location to the mouse, the extent of the trajectories were restricted as mice had limited space to initiate avoidance behavior. This limited space resulted in collisions with the obstacle and more time spent closer to the obstacle (S2 A,C). We therefore focused analysis on the distant obstacle location allowing enough space for mice to view the obstacle and update their steering. Therefore subsequent analysis incorporating the above criterion included about 25% of total trials in the light and dark, and 15% in the monocular condition (light: 2214/8881 trials, dark: 1456/5672 trials, monocular: 1274/8317). Results for the close location are included in the supplemental results (Fig S2).

### Vision guides directed paths around an obstacle at a distance

As nocturnal animals with low acuity vision, it is uncertain to what extent mice would use vision to avoid an obstacle at distance. To test the contribution of vision in our obstacle avoidance task, we compared trajectories of mice in light and dark conditions (see Methods). Under light conditions mice had overall more spatially efficient paths, spending less time closer to the obstacle (Fig 2B) and having overall less tortuous trajectories (Fig 2C), whereas in the dark a majority of trajectories went right up to the obstacle, indicating that mice contacted the obstacle before steering around it (Video S1). These data illustrate that mice can specifically utilize vision to help guide paths around an obstacle.

**Figure 2.**
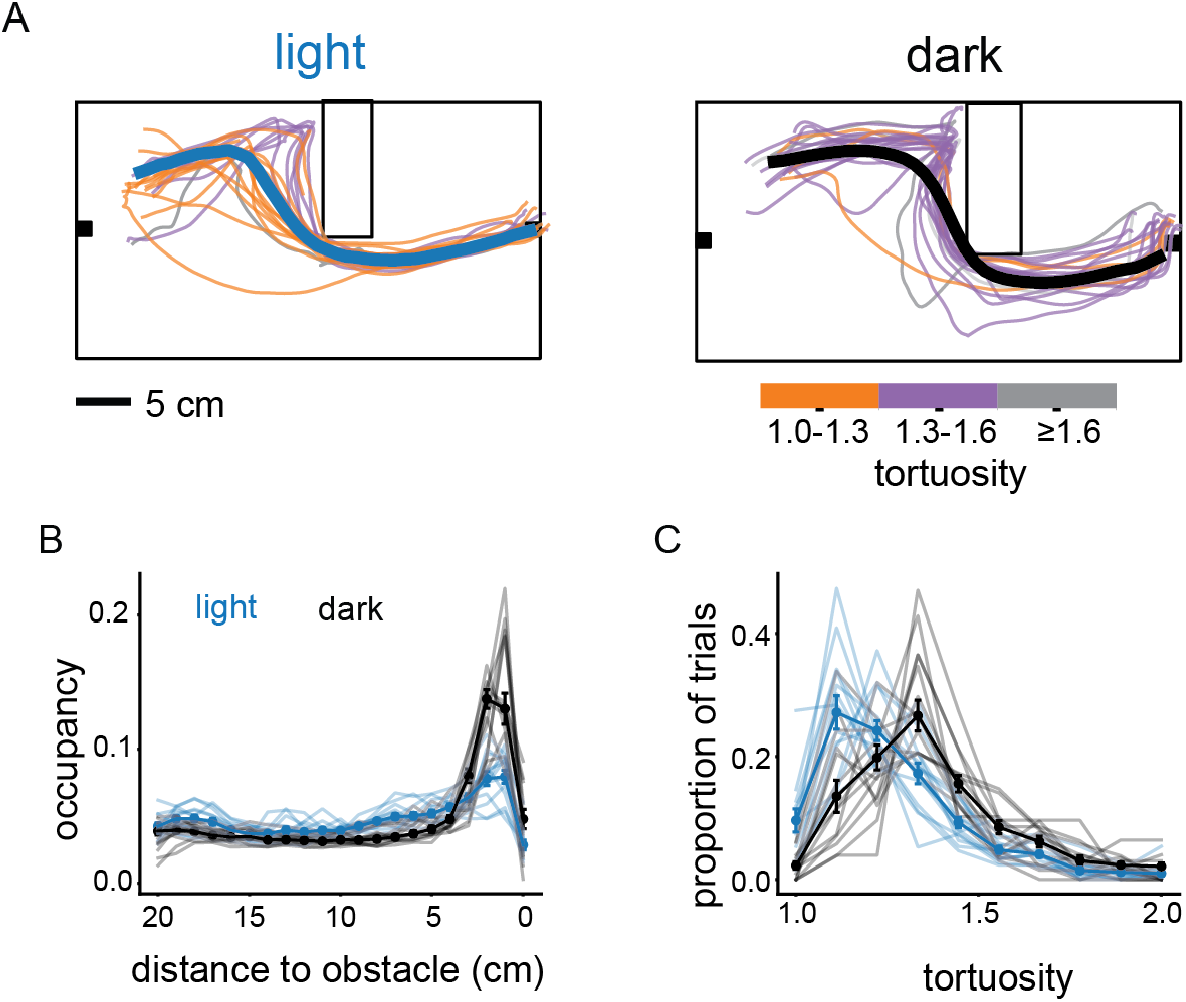
Role of vision in obstacle avoidance. (A) Twenty representative trajectories from a single mouse shaded based on tortuosity for one position of the obstacle in light and dark, population mean trajectory in bold. (B) Between-animal and single-animal median distribution of relative distance from an obstacle in light and dark. (C) Between-animal and single animal median distribution of tortuosity in light and dark. N = 13 mice.

To better understand how vision contributes to the behavioral dynamics of obstacle avoidance, we quantified how mice oriented and locomoted relative to the edge of the obstacle on the side of the open path (Fig 3A). We determined the angle of the mouse’s head relative to the obstacle edge, termed “heading to the opening”, and the lateral distance from the mouse’s head to the opening, termed “lateral error” (Fig 3B,C). When vision could be used, mice began orienting (Fig 3D) and moving (Fig 3E) to the opening starting at about 10 cm from the obstacle, well before they could use tactile information. On the other hand, in the dark, mice would generally continue at their initial heading and lateral error until they contacted the obstacle, then abruptly turn to and move towards the opening (Fig 3A,D,E,Video S1). Furthermore, in the light, mice also had more directed paths towards the goal ports compared to in the dark, demonstrating the need for vision in accurately targeting a known fixed location, in addition to avoiding an obstacle (Fig 3F,G). Taken together, this shows that mice use vision to avoid obstacles at a distance, directing heading and locomotion towards the edge of the obstacle closest to the open path.

**Figure 3.**
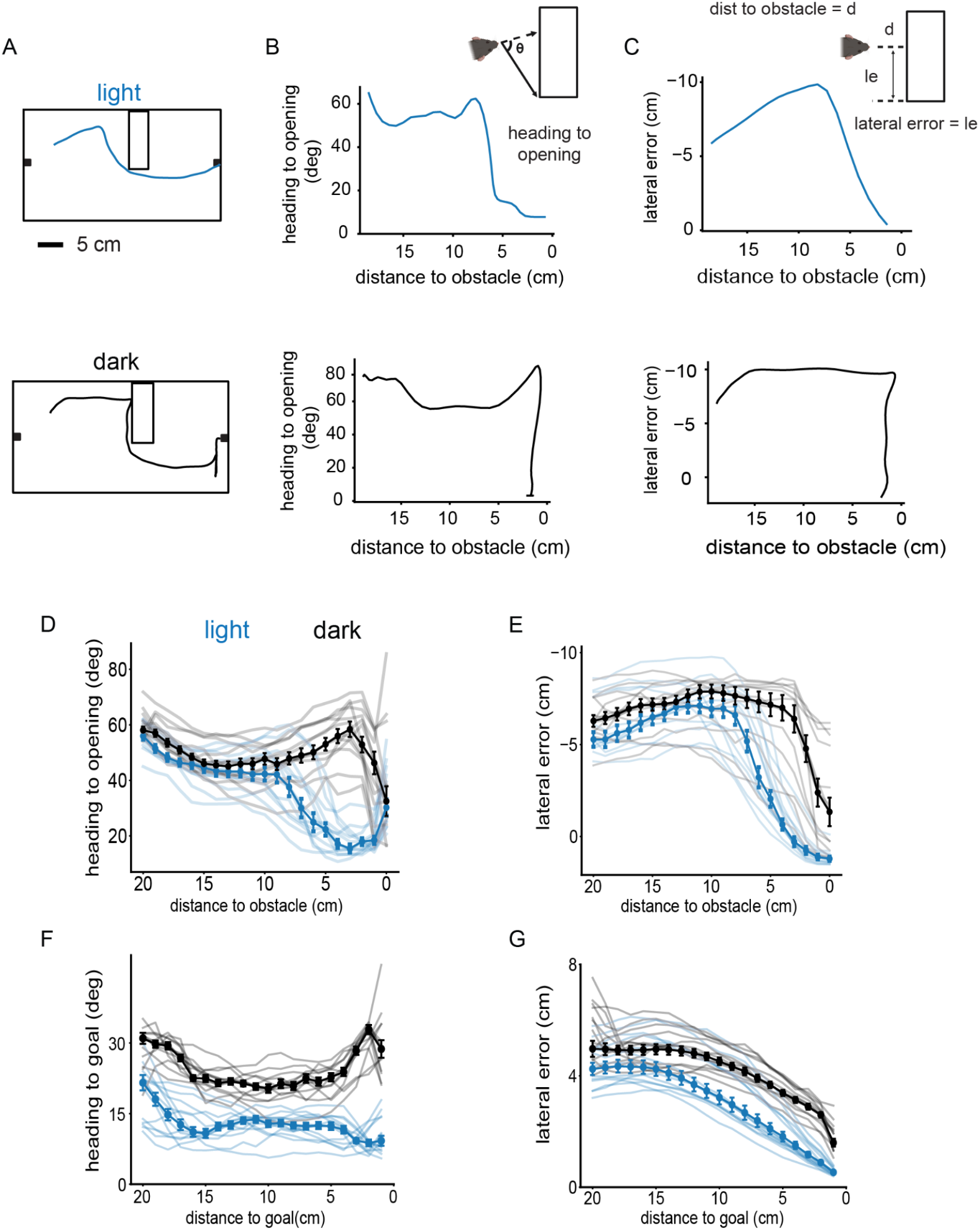
Vision guides directed steering around obstacles and approaches to goals (A) Example light and dark trajectories. (B-C) Corresponding heading to opening and lateral error as a function of distance to obstacle of example trajectories. (D-E) Between animal and single animal medians of heading to opening and lateral error. (F-G) Between animal and single animal medians of heading to goal and lateral error as a function of distance to goal. This included the portion of trajectories when the mouse had already passed the obstacle. N = 13 mice.

### Avoidance paths are guided by large orienting movements

To determine behavioral strategies for obstacle avoidance, we quantified the mouse’s turning based on the pattern of head movements before reaching the obstacle edge. We identified distinct head movements from peaks in head angular velocity, defined as peaks that had an absolute value >100 deg/sec and occurred >50 milliseconds apart. Within individual trials mice made large and abrupt turns, along with gradual turns via smaller head movements (Fig 4A,D).

**Figure 4.**
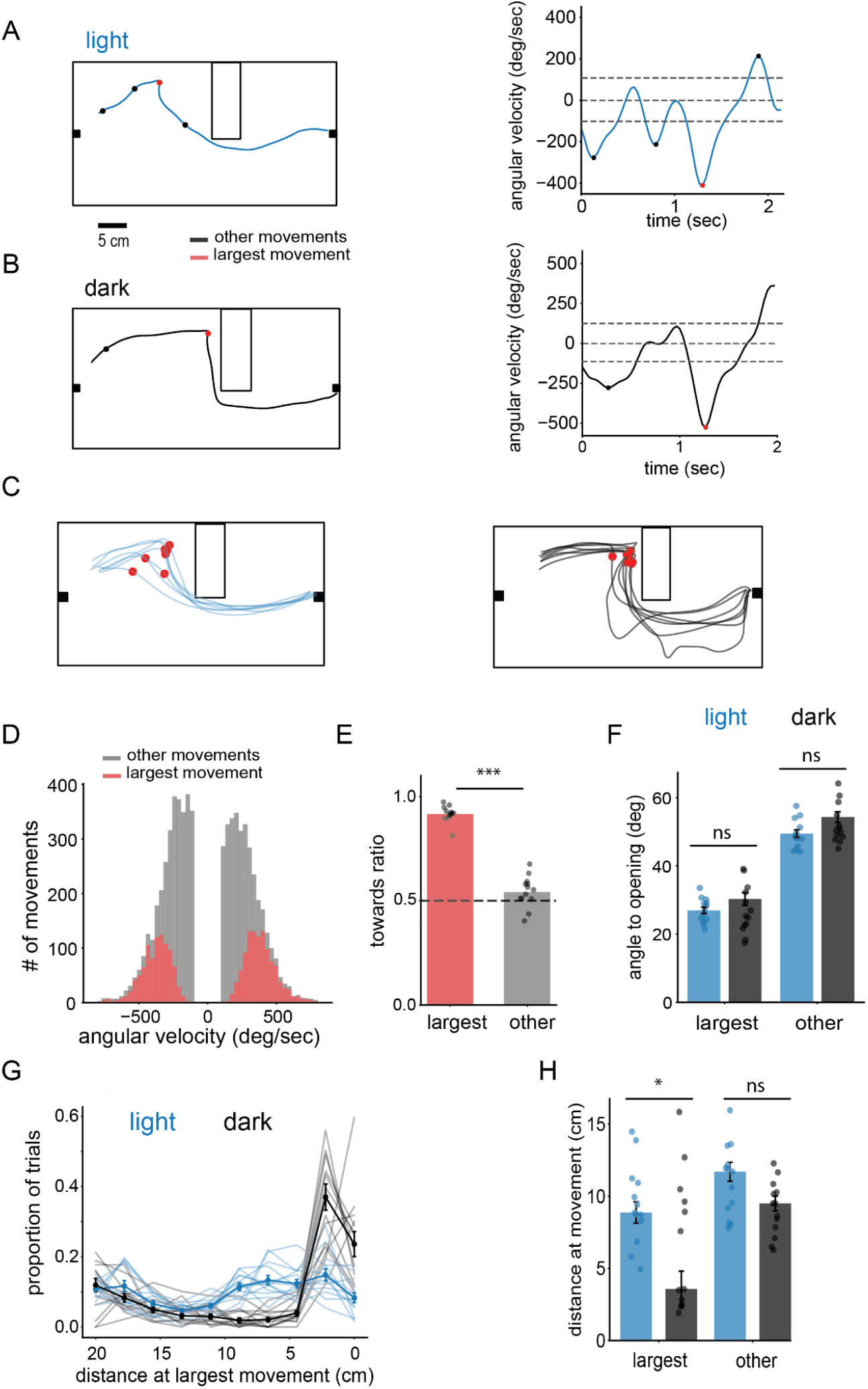
Vision facilitates directed head movements towards the obstacle opening at a distance (A-B) Left: example trajectory with location of head movements in light and dark. Right: corresponding angular velocity of example trajectory. Dotted line denotes 0 deg/sec and ± 100 deg/sec. (C) Twenty example trajectories with location of largest movement in red. (D) Population distribution of all head movements colored by movement category for light data. (E) Mean population towards ratio of largest and other head movements in the light. (F) Between-animal median of angle to opening for each movement category. (G) Between-animal median distribution of distance at largest movement with single animal medians. (H) Between-animal median of distance at movement for each movement type for dark and light. N =13 mice.

We then separately analyzed the largest movements within a trial from all other movements. Across light and dark conditions these large movements accounted for about 30 percent of all movements (Light: 31.2 ± 1.6 %, Dark: 32.9 ± 2.0%) and on average mice made 3 total movements (Light: 3.3 ± 0.2, Dark: 3.2 ± 0.2) per trial (Fig 4D, S3 A,C). By examining the direction of the head turn relative to location of the opening, we find that about 90 percent (Light:

91.7 ± 3.9 %, Dark 92.1 ± 4.0 %) of these large orienting movements were directed towards the opening, while smaller movements (Light: 54.5 ± 7.4 %, Dark 56.4 ± 9.6 %) were more evenly distributed between towards and away from the opening (Fig 4 E,S3B). Furthermore in the light and dark, heading towards the opening was much more accurate following the largest orienting movements (Light: 26.7 ±0.9°, Dark: 25.3 ± 1.9°) compared to all other orienting movements (Light: 48.4 ± 1.1°, Dark: 50.4 ± 1.5°) (Fig 4F). The magnitude of each movement category (Largest:Light: 455.7 ± 18.7°/sec, Dark: 494.7 ± 35.9°/sec, Other: Light: 253.3 ± 10.3°/sec, Dark: 261.8 ± 8.9°/sec) was also consistent across light and dark conditions (S3D).

We next examined the distance from the obstacle at which mice make these large orienting movements. In the light, mice make these large orienting movements starting at 10 cm from the obstacle’s edge, well before tactile information can be used (Fig 4C,G). In the dark these movements often occur very close to the obstacle edge, typically after contacting the obstacle (Fig 4C,G). Overall, these large edge-directed movements occur at much farther distances in the light than in the dark (Light: 8.9 ± 0.7 cm, Dark 3.6 ± 1.2 cm, P = 0.024; Wilcoxon signed rank) (Fig 4H). Other movements occurred at similar distances in light and dark. These findings clearly show that the large orienting movements we have identified are specifically directed at the obstacle edge. However, they occur at a distance in the light, suggesting they are using vision to identify the edge, while in the dark they generally result from hitting the obstacle, which provides information about its location and the direction to the edge.

### Mice can accurately avoid the obstacle without binocular cues

We next tested whether mice need binocular cues, such as disparity, or whether monocular cues are sufficient for obstacle avoidance. We sutured closed the left eyelid of eight mice that were previously trained in the obstacle avoidance task, eliminating binocular information. However, it is important to note that the visual field of each eye extends ∼20deg across the midline, so mice were still able to see directly in front of them with the non-occluded eye. Furthermore, the occlusion had only small impacts (<10 deg) on how mice kept their head oriented, and they kept their head aligned in the same direction they were moving irrespective of monocular occlusion (S4 G).

We first looked at the ratio of left to right turns out of the starting port, to see if mice would change their turning preference when monocularly occluded (Fig 5A). We found that mice tended to decrease their leftward preference in the monocular condition compared to their preference in the binocular condition (Binocular: 46.2 ± 5.6%, Monocular: 31.4 ± 7.9%, Paired t test; p = .38), consistent with a shift towards using the open-eye to view the arena upon exiting the port (Fig 5A). To account for the difference in visual information available depending on the direction the animals leave the port, we then split subsequent analysis into initial left and right turn trials. On left trials, the open side of the obstacle would be on the closed-eye side, while on right trials the obstacle opening would be on the open-eye side (Fig 5B).

**Figure 5.**
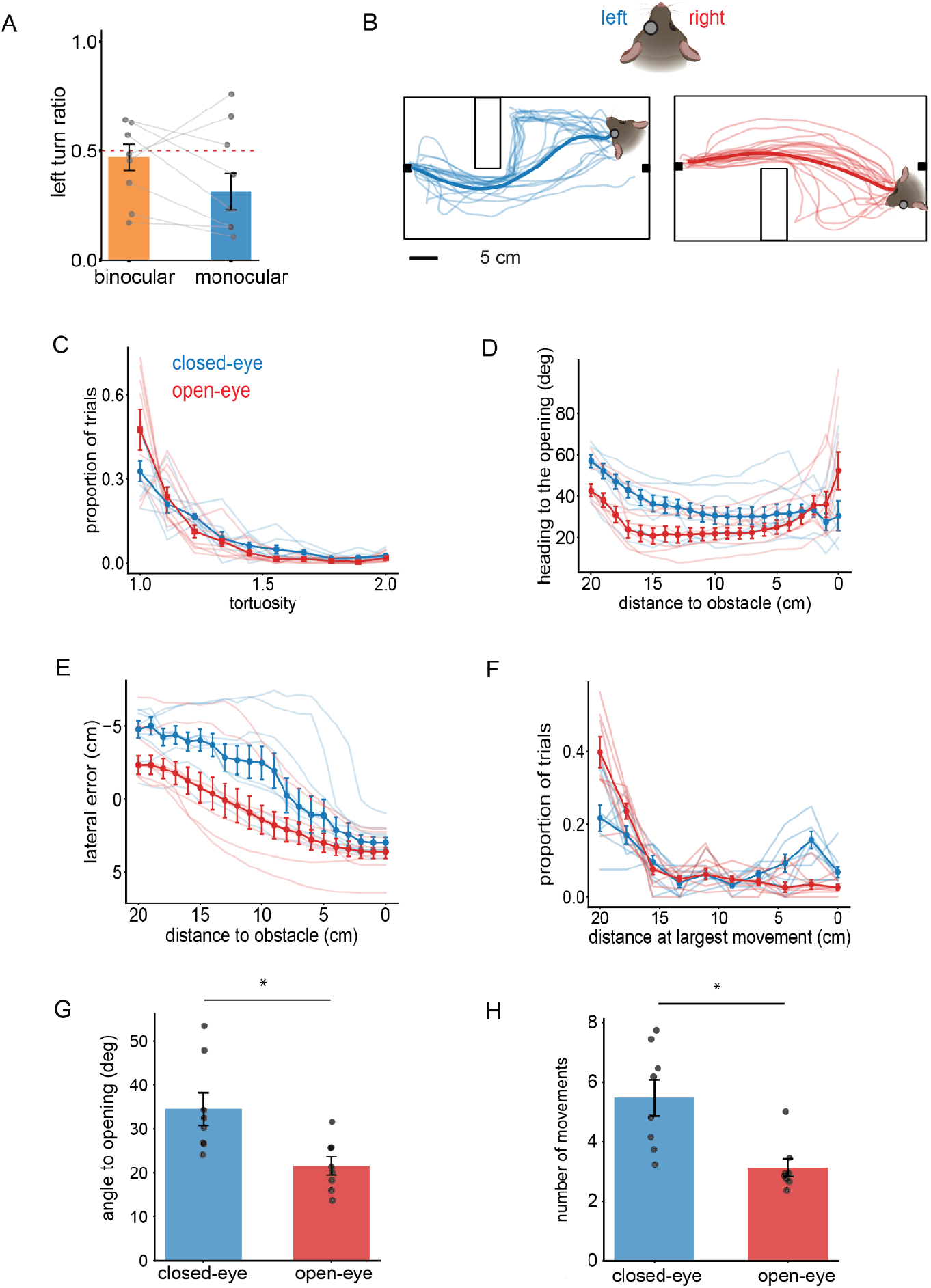
Monocular vision is sufficient for obstacle avoidance (A) Between-animal left turn ratio in binocular and monocular conditions. (B) Twenty example trajectories of left and right turns in monocular occluded mice with between animal median bolded.(C) Between-animal and single animal median of tortuosity of occluded and non-occluded side trials. (D-E) Between animal and single animal medians of heading to opening and lateral error as a function of distance to the obstacle edge of occluded and non-occluded side trials. (F) Between-animal and single animal median distribution of distance at largest movement relative to obstacle edge of occluded and non-occluded side trials. (G-H) Between-animal and single animal median of angle to opening following largest movement and number of total movements of occluded and non-occluded side trials. N=8 mice.

Trials leading with the closed-eye were more tortuous compared to binocular and open-eye trials, while open-eye trials had comparable tortuosities to binocular trials (Fig 5C, S4 A). Although closed-eye trials were more tortuous, they did not spend more time closer to the obstacle’s edge, relative to binocular trials, demonstrating they did not tend to hit the obstacle as seen in the dark condition (S4H). Trials where the animal led with the open-eye spent less time closer to the obstacle (S4H). These results suggest that limiting visual input via monocular sutures did not affect mice’s ability to avoid an obstacle. By observing the behavioral dynamics during obstacle avoidance, we found that on closed-eye trials mice were slightly less directed in their heading towards the opening (Fig 5D) but moved towards the opening at similar distances to binocular trials (Fig 5E, S4 C). Correspondingly, on open-eye trials mice moved and oriented towards the opening at farther distances relative to the obstacle edge compared to closed-eye and binocular trials (Fig 5D,E; Fig S4B,C).

Across both open-eye and closed-eye trials, about 90 percent of the largest orienting movements were directed to the opening (Closed-eye: 88.9 ± 7.4%, Open-eye: 93.5 ± 3.6%), similar to binocular conditions. On closed-eye trials there was a higher distribution of these large orienting movements closer to the obstacle compared to open-eye and binocular trials (Fig 5F, S4 D). On closed-eye trials, mice had similar accuracy in orienting to the opening as binocular trials (Fig 5G). Open-eye trials were significantly more accurate than closed-eye trials (Fig 5G). There was also an increase in the number of average total head movements per trial (Closed-eye: 5.49 ± 0.57 movements, Open-eye: 2.90 ± 0.27 turns, p = 0.039 ; Wilcoxon signed rank = 3.000) on the closed-eye trials compared to open-eye trials (Fig 5H), which may represent the need to acquire additional visual information.

This difference in directedness towards paths around the obstacle for the open-eye trials is likely due to the propensity of mice to move in the direction of the open-eye, i.e the area with visual information and the location of the opening. This is most evident in the difference in lateral error and heading to obstacle opening in the open-eye trials, as mice move to the opening sooner when it is visible to their open-eye (Fig 5 D,E). This further supports that in this task mice are actively using visual information, reacting to the obstacle while moving towards the goal. This was shown on closed-eye trials as mice made more head movements potentially to gain visual access to the obstacle opening (Fig 5 H). In summary, these data show that mice do not require binocular cues like stereopsis, and that monocular cues are sufficient for obstacle avoidance.

## Discussion

In the present study, we establish an obstacle avoidance task and demonstrate that mice use vision to avoid an obstacle at a distance, via discrete and directed orienting movements. Additionally, we show that mice can perform obstacle avoidance with monocular vision, making this paradigm amenable for studying the ethological capabilities of monocular vision in mice. Recent work using naturalistic visual-motor tasks have focused either on pursuit, escape behavior, or gap crossing^6^. Our demonstration of visually guided steering for obstacle avoidance provides additional insight into the repertoire of naturalistic visually guided behaviors in laboratory mice.

Quantification of behavioral dynamics and orienting movements revealed active visual control of steering during obstacle avoidance. Mice began to direct their heading and moved towards the opening edge simultaneously at about 10 cm from the obstacle. These paths are often guided by large orienting movements that result in accurately directed headings towards the opening. These results align well with previous findings demonstrating that mice and other rodents use vision to guide orienting under various behavioral contexts^1,4,20^. However, obstacle avoidance provides a very different visual context from other orienting behaviors, such as target pursuit. In both behavioral and physiological studies of pursuit behavior, researchers have shown that small and often moving visual stimuli drive approach^20,21,22^. Our obstacles are large and stationary, presenting very different visual stimuli than those reported for pursuit tasks. Our initial findings suggest that mice target the edge of the obstacle with large orienting movements^1,4^. It remains to be determined if other visual features of the obstacle besides its edge guide this orienting behavior or if the perception of the edge itself is sufficient for obstacle avoidance.

The results from monocular occlusion suggest that mice do not need visual cues that rely on binocular integration, like stereopsis to estimate distance, for detecting the obstacle and path around it. Patterns of visual motion or optic flow have been proposed as an important cue for guiding steering, and previous work shows that rodents use optic flow to guide deceleration in a collision task^23,24,25^. Recent work in humans and rodents shows that during walking and pursuit, gaze aligns with the direction of locomotion, resulting in patterns of optic flow useful for navigation^2,26^. It is still unknown how optic flow may contribute to the perception and avoidance of obstacles and to what extent it may guide steering. Analysis of the optic flow patterns that occurs during obstacle avoidance would give insight into how global motion signals may contribute to visually guided steering. In sum, further experiments are needed to elucidate what visual information is used to drive targeted orienting movements during obstacle avoidance, and how mice acquire this information behaviorally to guide their paths.

The superior colliculus (SC) is ideally suited to study visually guided orienting, as its layered organization directly links visual input to motor output^27,28,29,30^. The SC plays a role in both orienting to a target, and away from threats, both of which can be visually driven^31,21,32^. In the obstacle avoidance task, this could take the form of orienting toward the opening, or away from the obstacle itself. Direct targeting of head movements toward the obstacle edge suggests orienting toward this point. However, future work could investigate the role of two major behavioral and anatomically distinct pathways of SC related either to defensive or targeting movements via crossed and uncrossed pathways^33,34^.

Taken together, our study highlights the value of using naturalistic tasks to study active vision, as mice reliably use vision to avoid an obstacle, and further demonstrates that mice interact with their environment using vision. Because the ability to avoid obstacles is fundamental to survival, elucidating the mechanisms of this visually guided steering has the potential to provide important insight into conserved neural computation across mammals.

## Acknowledgements

We thank Dr. Jennifer Hoy, Dr. Phil Parker, and Niell lab members for feedback on the manuscript. This work was supported by NIH grants R01NS127305 (C.M.N.) and R01NS121919 (C.M.N.).

## Contributions

M.S and C.M.N. conceived the project. C.M.N. supervised all aspects of the project. M.S. led the mouse experiments. K.J. contributed to mouse experiments. M.S led data analysis. M.S and C.M.N. contributed to writing and editing the manuscript.

## Methods

### Animals

All procedures were conducted in accordance with National Institutes of Health guidelines and were approved by the University of Oregon Institutional Animal Care and Use Committee. 5-10 month-old mice (Mus musculus, C57BL/6J, Jackson Laboratories and bred-in house) were kept on a 12-h light/dark cycle. In total 12 male and 5 females were used for this study. Mice were housed with sibling cagemates for all conditions besides monocular deprivation experiments in which mice were singly housed. Humidity was between 40-60% and temperature was 21±1°C. Mice were placed under a water restriction schedule at the start of training, receiving their full daily water during training/task periods.

### Hardware and Arena

All experiments were done in a 50 × 25 cm rectangular arena with 40 cm high walls made of white acrylic and a gray silicone kitchen mat was used as the floor. The area was illuminated with visible light LEDs and infrared LED lights for video acquisition. Water was delivered using a 5V 3-way solenoid (Lee company LHDA0531115H) controlled by an IR photoelectric switch activated with nose pokes. These hardware interfaced with the acquisition computer using an Arduino UNO R3 with a 5V expansion shield (Gravity: IO Expansion Shield for Arduino V7.1). During the obstacle avoidance task a 13cm x 25cm x 5cm obstacle made of white acrylic was placed in the arena. Overhead video was taken at 60 fps with an infrared-sensitive FLIR BlackFly S USB3. For darkness experiments the housing apparatus was fitted with black out material (Thorlabs) and all room lights were turned off or covered. Timestamps were recorded from water delivery, nose pokes, and overhead camera using Bonsai (https://bonsai-rx.org/).

### Monocular suture

For all procedures, anesthesia was induced at 3% isoflurane and maintained at 1.5–2% in O2 at a 1 l/min flow rate. Ophthalmic ointment was applied to both eyes, and body temperature was maintained using a closed-loop heating pad at 37°C. The area immediately surrounding the eye to be sutured was wiped with 70% ethanol before ophthalmic ointment was applied. Two to three mattress sutures were placed using 6-0 silk suture, opposing the full extent of the lid. The forepaw and hindpaw nails ipsilateral to the sutured eye were trimmed to help minimize post-procedural injury.

### Preprocessing of trajectories

A DeepLabCut model (DLC 2.3.0) was used for post hoc keypoint tracking of the mouse and obstacle. The model was trained to track 8 points distributed between the nose and tail end of the mouse as well as the obstacle corners, arena corners, and water ports. All points were converted from pixels to cm. A trajectory was defined as the path of the nose point from the point of exit (5 cm away, facing forward) at the last port to the entry of the next active port.

### Behavioral metrics

*Tortuosity* for each trial trajectory was calculated by dividing the actual path length by the euclidean distance between the first and last point of the trajectory. *Heading to the obstacle/goal* was calculated from the angle between the vector of the neck to nose and neck to the corner adjacent to the openside or the center of the goal port. We quantified *lateral error* as the horizontal distance between the mouse’s nose and the corner of the openside or the center of the goal port. *Heading* was calculated as the angle measured between the neck and nose. We defined *head turns* as peaks in angular velocity that exceed 100 deg/s and occur at least 50 milliseconds from each other.

### Quantification and statistical analysis

Data were statistically analyzed using custom scripts written in Python (v.3.8, python.org). We used paired T tests for comparison of means and Wilcoxon signed rank for comparison of medians between conditions. Two-way ANOVA was used to compare the effects of different conditions. All error bars represent the standard error of the mean (SEM). In figures, * p < 0.05, **p <= 0.01, and ***p <= 0.001.

## Supplemental Figures

**Supplement 1.**
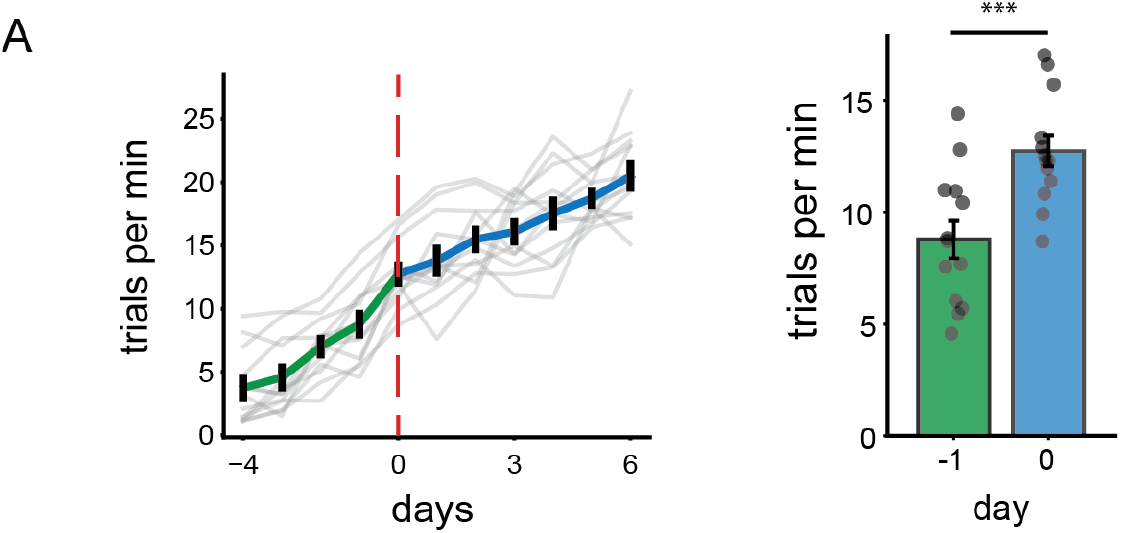
Obstacle avoidance task training (A) Left: Trials per minute of non-obstacle and obstacle avoidance. Right: Between animal and single animal mean trials per minute for the last day of non-obstacle and first day of obstacle (N=8).

**Supplement 2.**
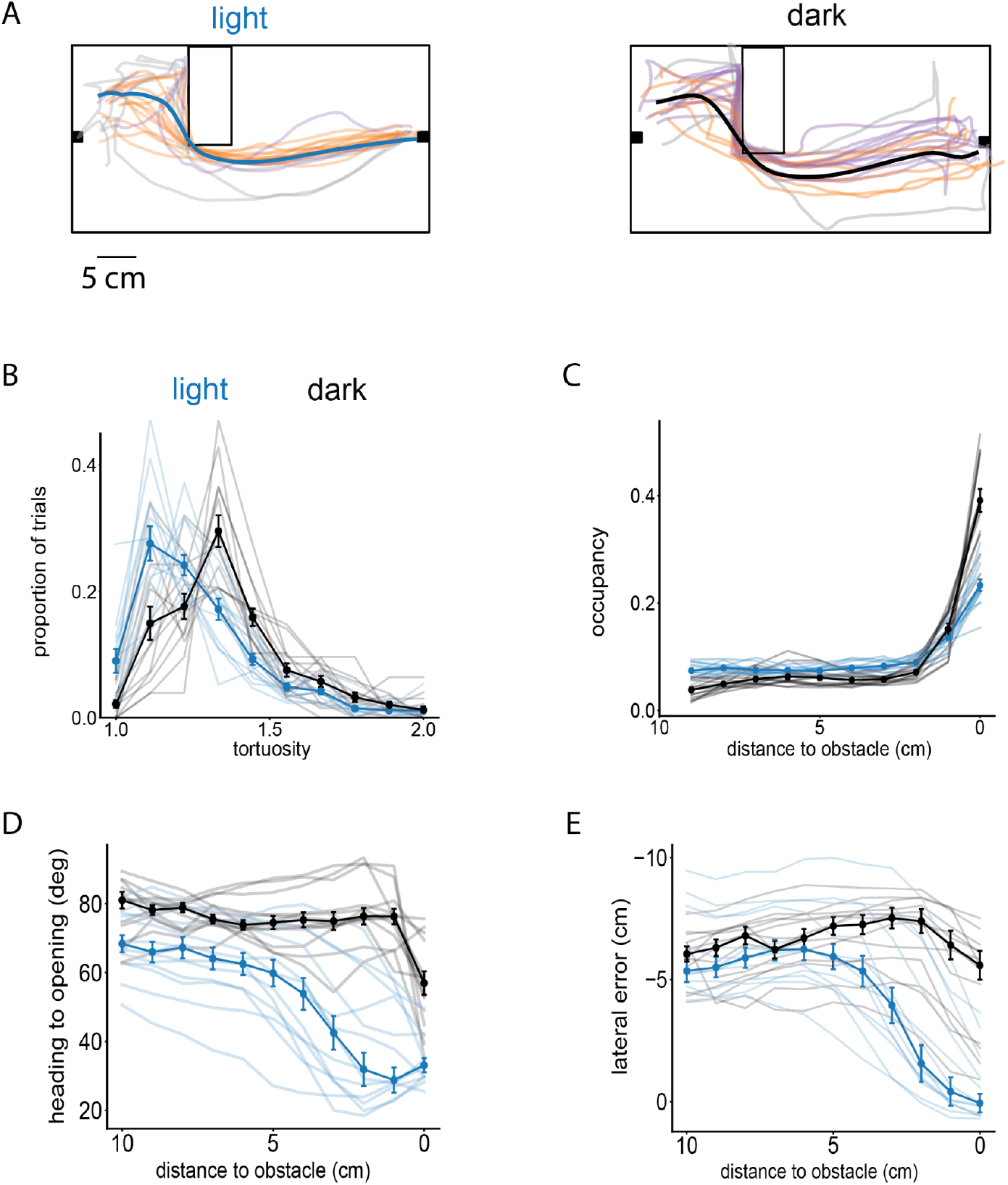
Obstacle avoidance behavior at short distances (A) 20 representative trajectories from a single mouse shaded based on tortuosity for one position of the obstacle in light and dark, population mean trajectory in bold. (B) Median population distribution of relative distance from an obstacle in light and dark. (C) Median population distribution of tortuosity in light and dark. (D-E) Population and single animal medians of heading to opening and lateral error. Single animal mean traces unbolded. N = 13

**Supplement 3.**
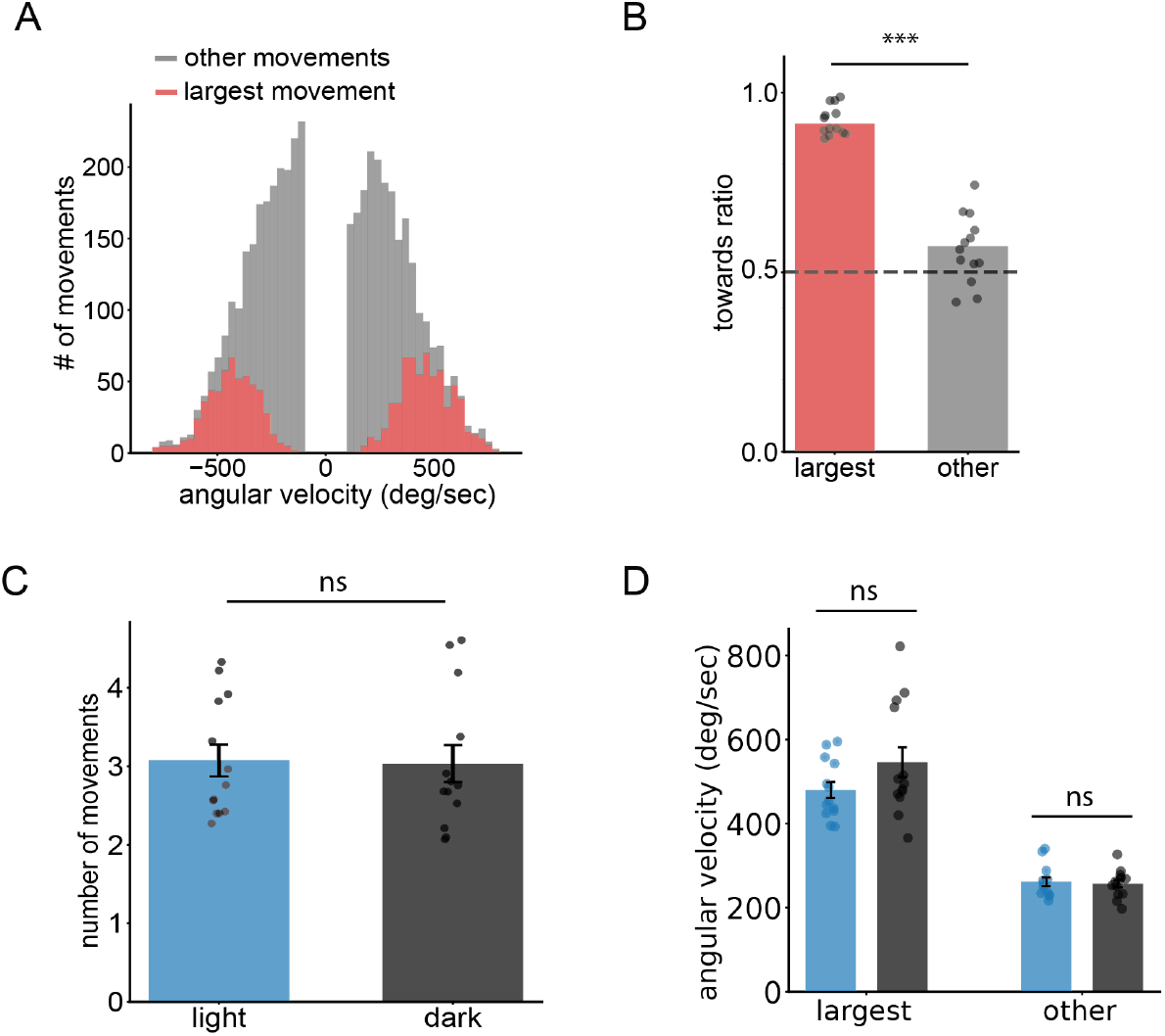
Additional head movement data (A) Population distribution of all head movements colored by movement category for dark data. B) Mean population towards ratio of largest and other head movements in the dark (C) Population median number of head movements per trial in light and dark. (D) Population median of angular velocity of largest and all other head movements. N = 13

**Supplement 4.**
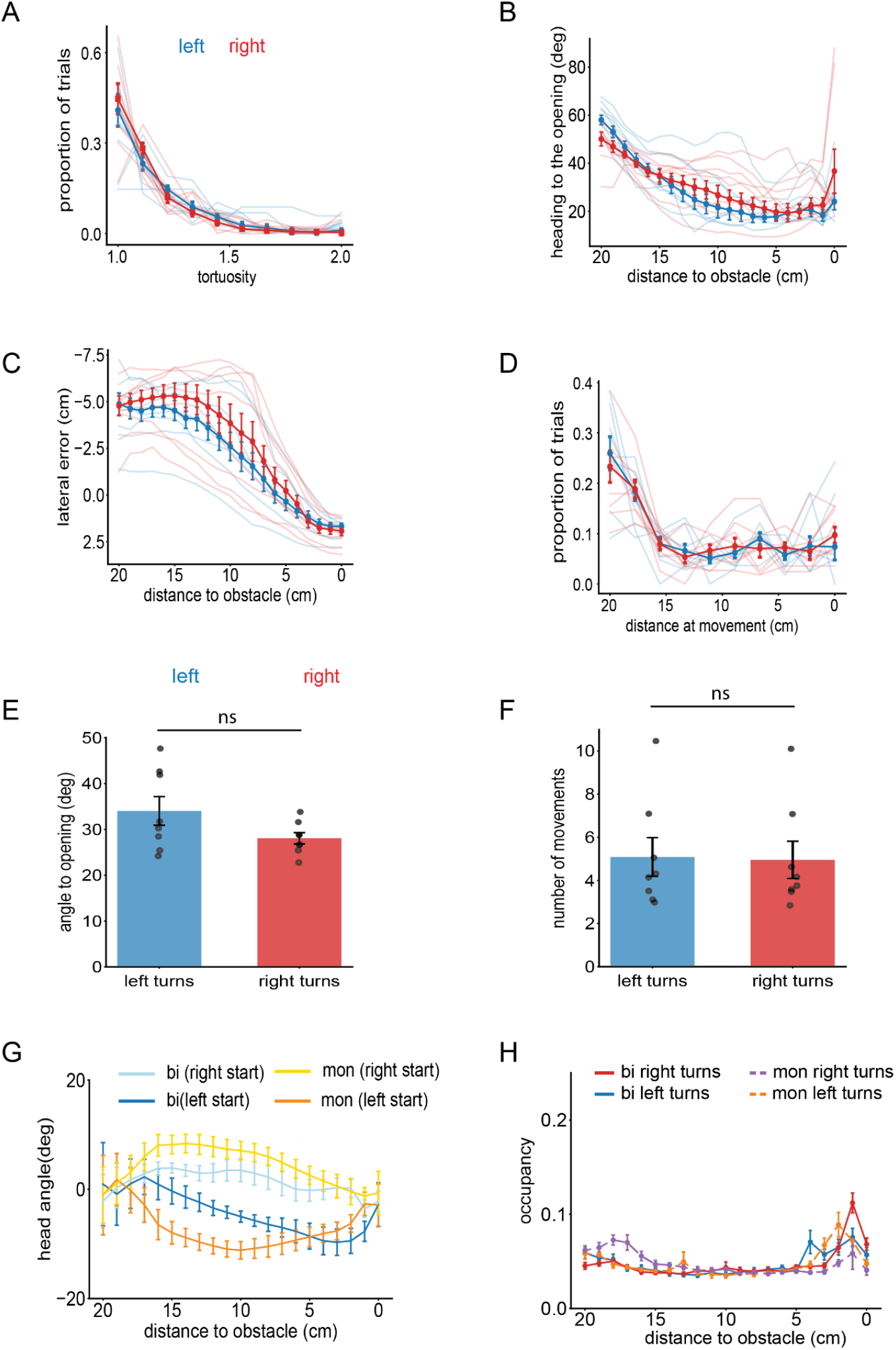
No difference in obstacle avoidance behavior across binocular left and right turns and additional monocular data. (A-B) Population medians of heading to opening and lateral error as a function of distance to the obstacle edge during binocular condition in left and right turn trials. (C) Median population distribution of tortuosity during left and right turn trials. (D) Median population distribution of distance at largest movement relative to obstacle edge in left and right turn trials. (E-F) Population median of angle to opening following largest movement and number of total movements in left and right trials.(G) Between animal median of head angle relative to distance to obstacle edge. (H) Between animal median distribution of relative distance from the edge. N=8 mice.

**Supplement 5.**
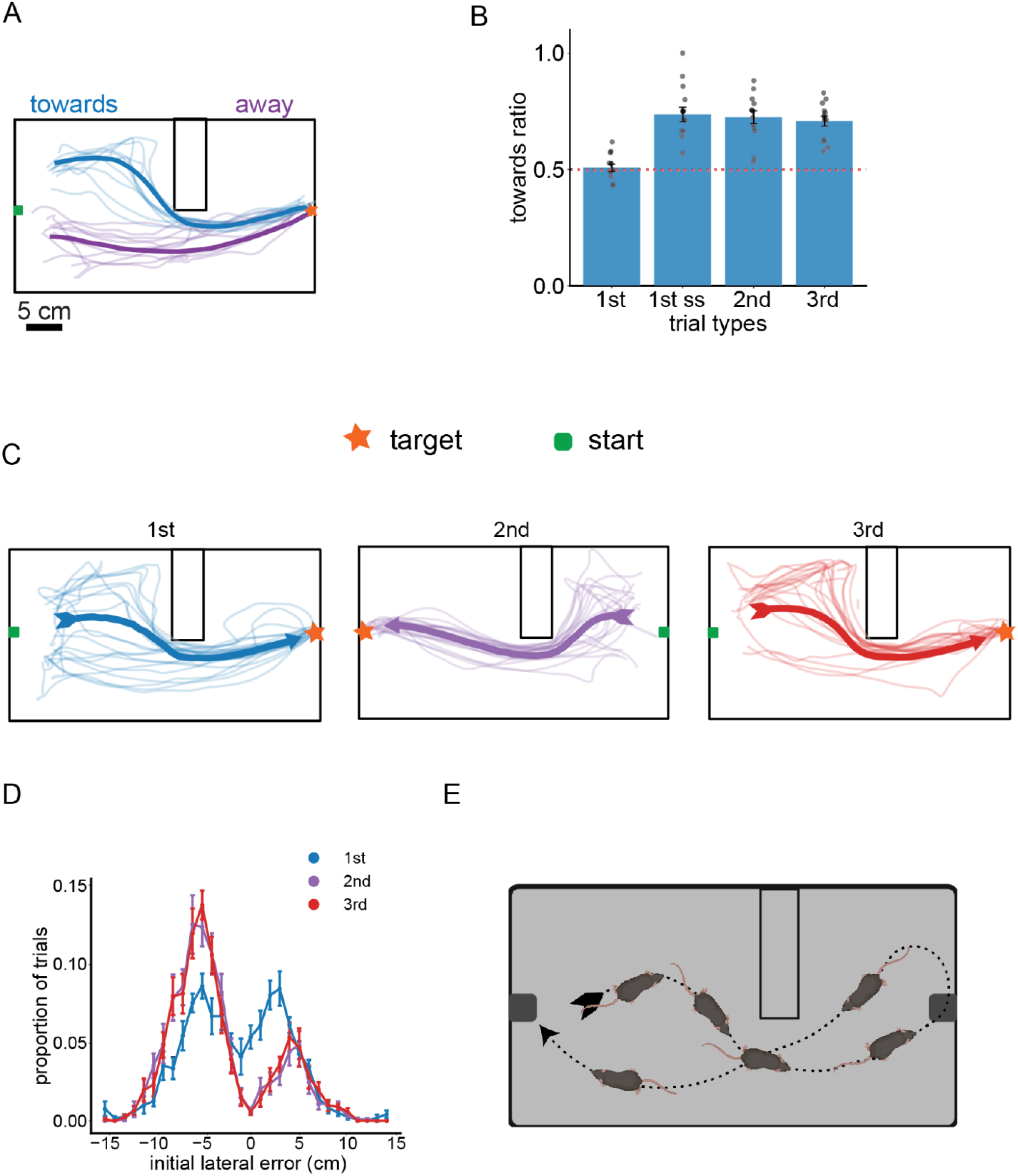
Effect of repeated presentations of obstacle location on steering behavior (A) Between animal mean and example trajectories of towards and away trials for a single obstacle position. (B) Towards ratio at different number of presentations of the obstacle at the same location. SS = same side, i.e. in the previous trial the obstacle was on the same side but moved to either the far or close location. (C) Between animal mean and single trajectories for corresponding trial type at a single obstacle location. (D) Between animal mean distribution of initial lateral error for corresponding trial type. (E) Cartoon depiction of standard mouse trajectories, demonstrating why mice tend to turn towards the obstacle despite seeing the obstacle at the same location on previous trials, due to circling behavior.

